# Machine learning-based decoding of emotional valence from electrophysiological signals in the monkey brain

**DOI:** 10.64898/2026.05.13.724152

**Authors:** Nakamura Shinya, Tuo Xiaoying, Watanabe Hidenori, Takuya Sasaki, Ken-Ichiro Tsutsui

## Abstract

Understanding how the brain operates in naturalistic settings requires methods that go beyond conventional repeated-measurement approaches, necessitating the development of single-trial neural activity analysis. Recent advances in machine learning offer new opportunities for analyzing brain electrophysiological signals. Here, we recorded surface electrocorticography (ECoG) and intracranial local field potentials (LFPs) from emotion-related brain regions in a monkey performing a Pavlovian conditioning task, in which sensory cues predicting reward or punishment were presented randomly, followed by the actual unconditioned outcome. We evaluated the performance of two machine learning algorithms, a Convolutional neural network (CNN) model and a Transformer-based model (EEG-Conformer), in classifying raw ECoG/LFP traces. Both models successfully classified valence type during conditioned and unconditioned stimulus presentation. Furthermore, the Transformer achieved significantly superior classification performance compared to the CNN, particularly in multi-state classification including baseline periods. By optimizing the training dataset for the Transformer model, we could detect dynamic fluctuations in emotional valence consistent with task type from continuously evolving ECoG/LFP patterns recorded throughout the task. These results demonstrate the utility of Transformer-based models for decoding emotional valence from neurophysiological signals in non-human primates.

## INTRODUCTION

Recent behavioral and cognitive neuroscience has increasingly turned its attention to understanding how the brain operates in naturalistic and real-world settings, beyond findings obtained under controlled laboratory conditions ^1,2^. Under laboratory conditions, measurements are typically repeated under controlled conditions, with relatively low variability that allows additive analyses. In real-world settings, however, researchers often have to deal with single events that do not recur. Therefore, in addition to conventional approaches that integrate and analyze data obtained from repeated measurements, it is important to develop methods for analyzing and evaluating neural activity on a single-trial basis ^3^. Deep learning approaches have improved the decoding of neural signals ^4^.Transformer-based models demonstrated speech decoding from electrocorticogram (ECoG) signals that require real-time processing ^5-7^.

In our previous study, we reported that the Transformer neural network model is effective in discriminating local field potential (LFP) patterns, and that its discriminative power is higher than that of AlexNet-based models or simple LFP power analysis ^8^. The Transformer model could be trained to discriminate multi-channel LFP data recorded from mice experiencing abdominal pain induced by intraperitoneal acetic acid injection from data recorded under no-pain conditions, with high precision (*d*-prime = 3.37). For emotion classification, deep neural network models ^9^ have been proposed from publicly available datasets such as DEAP ^10^ or SEED ^9^. However, estimating emotional states from single-trial EEG signals remains challenging due to the low signal-to-noise ratio and substantial inter-subject variability in emotional responses. Direct recordings of neural activity from the cerebral cortex or deep brain regions are therefore required for fundamental investigations of real-time emotional decoding.

The purpose of this study was to further evaluate the validity of the Transformer model ^11^ in discriminating positive and negative emotional valence elicited in a more naturalistic manner in a monkey. We trained a monkey to perform a Pavlovian conditioning task in which syringes of different colors served as conditioned stimuli for forced liquid delivery into the mouth (fruit juice as a reward or saline as a punishment). Multiple electrocorticogram (ECoG) from the brain regions reflecting emotional states, which were the prefrontal ^12,13^, premotor ^14^, primary motor ^15^, and primary somatosensory cortices (PFC, PMC, MI and SI), as well as deep LFP signals from dorsal and ventral anterior cingulate cortex ^16^, amygdala ^17^ and nucleus accumbens ^18^ (dACC, vACC, Amy and NAc) were recorded and analyzed offline and demonstrated single-trial based decoding using a Transformer model.

## Results

### Pavlovian conditioning and brain ECoG/LFP recordings in the monkey

To induce positive or negative emotional valence in a monkey, we subjected a monkey to Pavlovian conditioning task following electrode implantation surgery mainly targeting dorsal and medial frontal cortices as well as nucleus accumbens and amygdala. In this task, a single experimenter presented a head-fixed monkey with differently colored syringes (orange or blue) containing either apple juice (reward) or saturated saline (punishment) at a distance of approximately 30 cm from the monkey’s face (Fig. 1A), after which the experimenter brought the tip of the syringe to the monkey’s mouth to allow licking so that the monkey voluntarily obtained the liquid from the syringe tip. In each trial, following a 5-s baseline (BL) period, a syringe containing either juice or saline was presented pseudo-randomly for 5 s; this period was defined as the conditioned stimulus (CS) period for reward and punishment, respectively (CSr and CSp) (Fig. 1B). Subsequently, the syringe was brought to the monkey’s mouth, and the monkey obtained the liquid for 5 s; this period was defined as the unconditioned stimulus (US) period for reward and punishment, respectively (USr and USp). The timing of syringe movements was controlled by an LED indicator positioned behind the monkey, which served as a timer to guide the experimenter’s actions. Following habituation to the experimental setup, in each experimental day, before starting the recordings, several training trials were performed to allow the monkey to learn the association between the syringe colors and the outcomes. After sufficient training, the monkey reliably licked during juice delivery (USr) but rarely licked during saline delivery (USp) (Fig. 1C), indicating that the monkey discriminated between syringe colors and generated positive or negative emotional responses accordingly. Following sufficient training, we analyzed results from a total of five recording days in which electrophysiological recordings were successfully performed.

**Figure 1.**
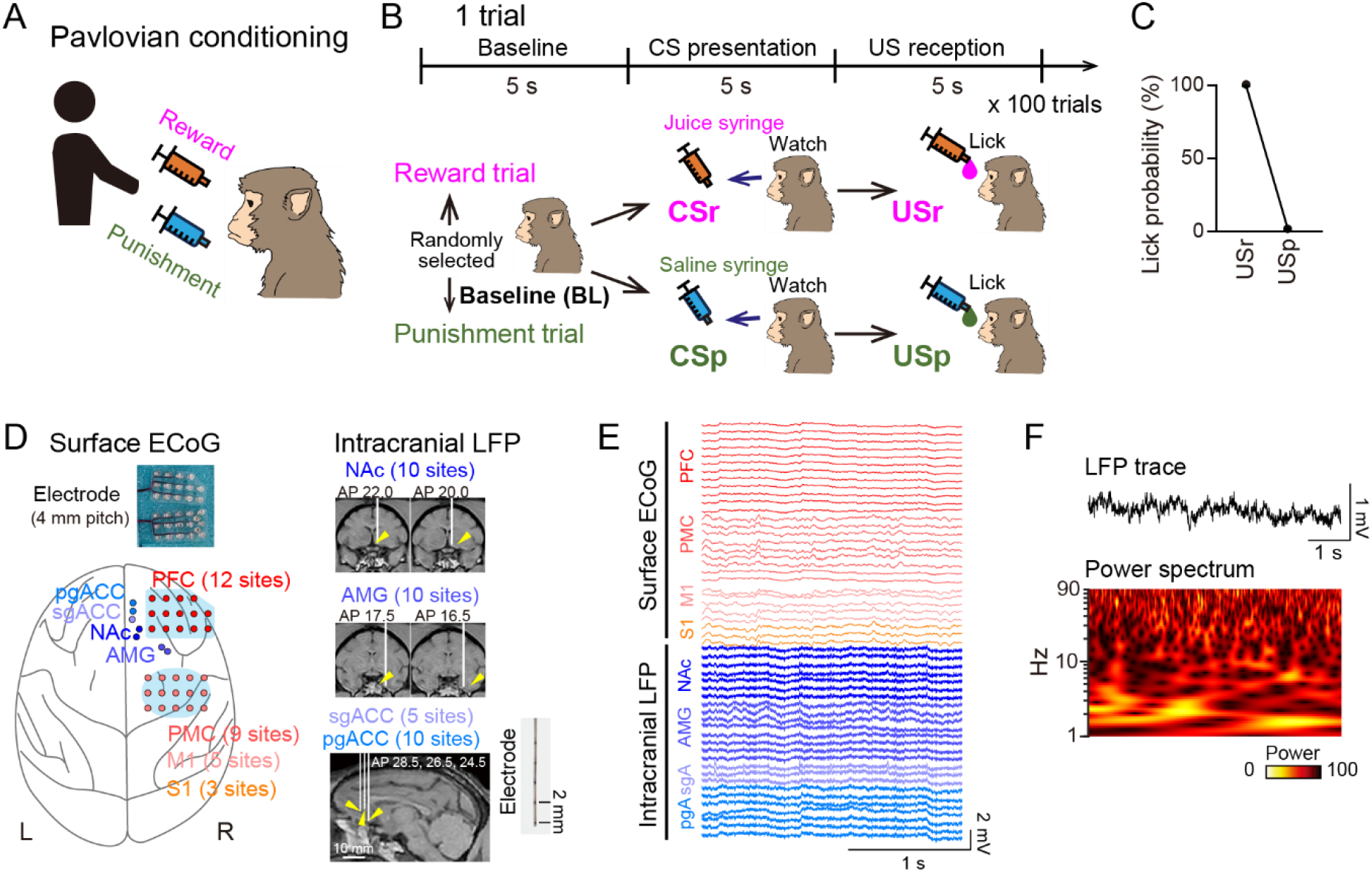
ECoG/LFP recordings from the monkey brain associated with emotional valence. (A) A well-trained, head-fixed monkey is presented with a colored syringe containing either juice (reward) or saline (punishment) by the experimenter. ECoG/LFP signals were monitored. (B) Trial structure. Each trial is randomly assigned as either a reward or punishment trial and consists of three consecutive 5-s periods: baseline (BL), conditioned stimulus (CSr or CSp) (syringe presentation), and unconditioned stimulus (USr or USp) (solution delivery with licking). (C) Licking probability during USr and USp periods (*n* = 453 and 454 trials from 5 days). Error bar represents SEM. (D) Surface ECoG and intracranial LFP recording locations. Yellow arrowheads indicate the position of electrode tips in the MRI image. (E) Representative ECoG/LFP signals from 54 recording sites. (F) A LFP wavelet spectrum in a trial.

In the monkey, two custom-designed ECoG electrode arrays (14 and 15 recording sites, respectively; 4 mm inter-electrode distance) were implanted to cover the cortical surface of the prefrontal cortex (PFC; 12 sites), premotor cortex (PMC; 9 sites), primary motor cortex (M1; 5 sites), and primary somatosensory cortex (S1; 3 sites) (Fig. 1D; surface ECoG). In addition, intracranial LFP electrodes with five vertically arranged recording sites (2 mm inter-electrode distance) were inserted into the nucleus accumbens (NAc; 10 sites), amygdala (AMG; 10 sites), subgenual anterior cingulate cortex (sgACC; 5 sites), and pregenual anterior cingulate cortex (pgACC; 10 sites). These electrode positions were verified by CT imaging. Electrophysiological recordings were performed throughout the task session. Electrophysiological recordings were performed throughout the task session. Typical ECoG/LFP traces simultaneously recorded from these regions are shown in Fig. 1E. These traces were first examined using wavelet power analysis (Fig. 1F).

### Frequency-power-based discrimination of reward and punishment

The averaged ECoG/LFP power was examined across six frequency bands: delta (1–4 Hz), theta (4–8 Hz), alpha (8–14 Hz), beta (14–30 Hz), slow gamma (30–80 Hz), and fast gamma (80–95 Hz) (Fig. 2A; *n* = 832, 393, 438, 392, and 444 trials from 5 recording days). For several electrodes showing statistically significant differences across the five periods (*P* < 0.05, Tukey’s test), the key features were as follows. Delta power was higher during the CSp period than during other periods in most electrodes, regardless of whether the recordings were obtained from surface ECoG or intracranial LFP. In particular, for surface ECoG electrodes, delta power during the CSp period was significantly greater than during the CSr period (*P* < 0.05), indicating that delta power more prominently increases during the CS period anticipating punishment. For theta power, the pairs of periods showing significant differences were not consistent across brain regions; however, theta power tended to be generally higher during CS periods than during US periods. Significant differences between CS and US periods were most frequently observed in PMC, M1, S1, and AMG (*P* < 0.05), suggesting that theta power is elevated during the presentation of conditioned stimuli (CS periods) compared with the US period during which outcomes are actually delivered. Alpha and beta power showed fewer significant period pairs; however, the overall trend was similar to that observed for theta power. In contrast, no significant differences were observed across the five periods for either low or high gamma power.

**Figure 2.**
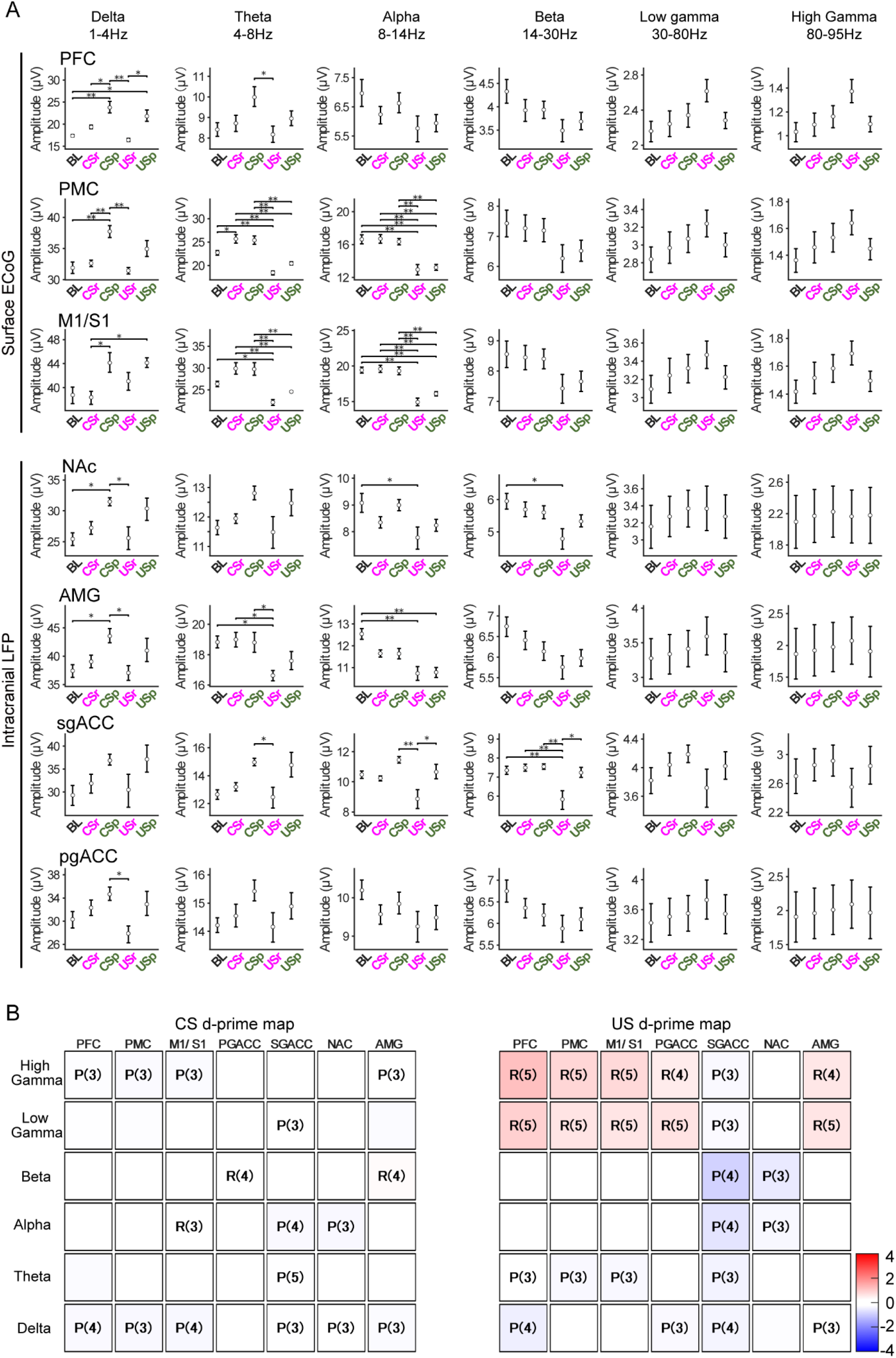
Frequency-band power across task epochs and conditions. (A) ECoG/LFP power in each region during individual five periods. Error bars represent SEM. *\*P* < 0.05, \**\*P* < 0.01, one-way ANOVA followed by Tukey’s post-hoc test. (B) Frequency-power-based discrimination of reward and punishment. Each box corresponds to a specific brain region (x-axis) and frequency band (y-axis). The color indicates the mean signed *d*-prime values across sessions; positive values indicate greater activity for reward than punishment, whereas negative values indicate the opposite. Letters indicate conditions with significantly larger baseline-corrected band-limited power in more than half of the total sessions (R: reward-dominant; P: punishment-dominant). Blank boxes did not meet this criterion. Quantitative values are provided in Table 1.

To further evaluate whether these power differences were sufficient to distinguish reward from punishment, we quantified reward/punishment separation using signed *d*-prime values calculated from trial-wise baseline-subtracted band-limited power. This analysis was performed because significant differences in averaged power across task periods do not necessarily indicate that reward and punishment conditions can be separated on a trial-by-trial basis within a given epoch. For each session, signed *d*-prime values were computed separately for the CS and US epochs in each brain region and frequency band (Fig. 2B and Table 1; n = 5 recording days). Across the examined brain regions and frequency bands, mean *d*-prime values were generally small, indicating that band-limited power provided only limited separation between reward and punishment and was not sufficient for reliable trial-by-trial discrimination. To visualize the directionality of trial-level effects, we further summarized whether reward or punishment trials showed significantly larger baseline-corrected band-limited power within each session. This analysis revealed only limited and region/frequency-dependent reward-or punishment-dominant patterns, with few combinations showing dominance in more than half of the recording sessions.

**Table 1.** Summary of frequency-power-based reward–punishment discrimination. This table provides the quantitative values underlying Figure 4. For each brain region and frequency band, mean *d*’ across sessions, and dominant condition are shown. Dominance is expressed as R or P (nR:nP), where nR and nP indicate the numbers of sessions classified as reward- or punishment-dominant, respectively. Cases not exceeding half of the total number of sessions are indicated by “−”.

| Epoch Type | Region | Band | Mean $d'$ | Dominant Condition (nR:nP) | Epoch Type | Region | Band | Mean $d'$ | Dominant Condition (nR:nP) |
| --- | --- | --- | --- | --- | --- | --- | --- | --- | --- |
| CS | PFC | $\delta$ | -0.332 | P (0:4) | US | PFC | $\delta$ | -0.482 | P (0:4) |
| | | $\theta$ | -0.294 | - (0:2) | | | $\theta$ | -0.217 | P (1:3) |
| | | $\alpha$ | -0.127 | - (0:0) | | | $\alpha$ | -0.110 | - (0:0) |
| | | $\beta$ | 0.038 | - (1:1) | | | $\beta$ | -0.211 | - (0:2) |
| | | Low $\gamma$ | -0.277 | - (0:2) | | | Low $\gamma$ | 0.928 | R (5:0) |
| | | High $\gamma$ | -0.288 | P (0:3) | | | High $\gamma$ | 1.165 | R (5:0) |
| | PMC | $\delta$ | -0.347 | P (0:3) | | PMC | $\delta$ | -0.243 | - (0:2) |
| | | $\theta$ | 0.054 | - (2:0) | | | $\theta$ | -0.339 | P (0:3) |
| | | $\alpha$ | 0.164 | - (2:0) | | | $\alpha$ | -0.007 | - (1:1) |
| | | $\beta$ | 0.139 | - (1:0) | | | $\beta$ | -0.210 | - (1:2) |
| | | Low $\gamma$ | -0.293 | - (0:1) | | | Low $\gamma$ | 0.639 | R (5:0) |
| | | High $\gamma$ | -0.313 | P (0:3) | | | High $\gamma$ | 0.847 | R (5:0) |
| | M1/S1 | $\delta$ | -0.358 | P (0:4) | | M1/S1 | $\delta$ | -0.197 | - (0:2) |
| | | $\theta$ | 0.040 | - (1:0) | | | $\theta$ | -0.337 | P (0:3) |
| | | $\alpha$ | 0.084 | R (3:1) | | | $\alpha$ | -0.236 | - (1:2) |
| | | $\beta$ | 0.088 | - (1:0) | | | $\beta$ | -0.174 | - (1:2) |
| | | Low $\gamma$ | -0.178 | - (0:1) | | | Low $\gamma$ | 0.618 | R (5:0) |
| | | High $\gamma$ | -0.306 | P (0:3) | | | High $\gamma$ | 0.804 | R (5:0) |
| | PGACC | $\delta$ | -0.095 | - (0:1) | | PGACC | $\delta$ | -0.325 | P (0:3) |
| | | $\theta$ | -0.155 | - (0:0) | | | $\theta$ | -0.124 | - (1:1) |
| | | $\alpha$ | -0.178 | - (0:1) | | | $\alpha$ | -0.127 | - (1:2) |
| | | $\beta$ | 0.226 | R (4:0) | | | $\beta$ | -0.098 | - (0:1) |
| | | Low $\gamma$ | -0.137 | - (0:1) | | | Low $\gamma$ | 0.649 | R (5:0) |
| | | High $\gamma$ | -0.243 | - (0:1) | | | High $\gamma$ | 0.524 | R (4:0) |
| | SGACC | $\delta$ | -0.275 | P (0:3) | | SGACC | $\delta$ | -0.400 | P (0:4) |
| | | $\theta$ | -0.277 | P (0:5) | | | $\theta$ | -0.414 | P (0:3) |
| | | $\alpha$ | -0.326 | P (0:4) | | | $\alpha$ | -0.635 | P (0:4) |
| | | $\beta$ | -0.022 | - (1:1) | | | $\beta$ | -0.903 | P (0:4) |
| | | Low $\gamma$ | -0.119 | P (1:3) | | | Low $\gamma$ | -0.319 | P (1:3) |
| | | High $\gamma$ | -0.006 | - (1:0) | | | High $\gamma$ | -0.333 | P (0:3) |
| | NAC | $\delta$ | -0.239 | P (0:3) | | NAC | $\delta$ | -0.284 | - (0:2) |
| | | $\theta$ | -0.140 | - (0:0) | | | $\theta$ | -0.199 | - (0:1) |
| | | $\alpha$ | -0.346 | P (0:3) | | | $\alpha$ | -0.365 | P (1:3) |
| | | $\beta$ | 0.128 | - (2:1) | | | $\beta$ | -0.556 | P (0:3) |
| | | Low $\gamma$ | -0.248 | - (0:2) | | | Low $\gamma$ | 0.215 | - (2:0) |
| | | High $\gamma$ | -0.111 | - (0:2) | | | High $\gamma$ | -0.018 | - (1:0) |
| | AMG | $\delta$ | -0.298 | P (0:3) | | AMG | $\delta$ | -0.249 | P (0:3) |
| | | $\theta$ | 0.042 | - (1:0) | | | $\theta$ | -0.219 | - (0:2) |
| | | $\alpha$ | -0.025 | - (0:1) | | | $\alpha$ | -0.080 | - (0:1) |
| | | $\beta$ | 0.305 | R (4:0) | | | $\beta$ | -0.185 | - (0:2) |
| | | Low $\gamma$ | -0.301 | - (0:2) | | | Low $\gamma$ | 0.622 | R (5:0) |
| | | High $\gamma$ | -0.261 | P (0:3) | | | High $\gamma$ | 0.593 | R (4:0) |

Although robust separation was not observed, *d*-prime values tended to be larger during the US epoch than during the CS epoch in several regions and frequency bands, suggesting that neural activity during the US period may contain somewhat stronger reward– punishment-related information than cue-period activity. Together, these findings suggest that frequency-band power, computed within each epoch without preserving within-epoch temporal structure, provided limited separation between reward and punishment. This motivated end-to-end decoding of minimally preprocessed waveforms.

### Classification of ECoG/LFP traces by machine learning

Next, we investigated whether machine learning approaches could more effectively discriminate valence-related ECoG/LFP patterns. We compared the classification performance of two architectures: a convolutional neural network (CNN) and the EEG-Conformer (CNN+Transformer) model (Fig. 3A) ^11^. Raw LFP traces were used as input without conversion to power spectra. Traces were downsampled to 200 Hz, yielding a 50 (recording sites) × 1000-dimensional tensor for each 5-s period (Fig. 3B). In each model, classification performance was evaluated using stratified ten-fold cross-validation, maintaining a train-to-test ratio of 9:1, in which 90% of trials were used for training and the remaining 10% for testing. As an initial assessment, we evaluated whether ECoG/LFP patterns could be classified between CSr and CSp periods (Fig. 3C), which do not involve direct sensory input or overt behavioral responses sas in the US period, but rather represent an anticipatory phase during which the monkey predicts the upcoming gustatory outcome. Models were trained to discriminate between CSr and CSp periods, and classification performance was assessed by constructing a confusion matrix and computing a *d*-prime value from each matrix. In all five recording days, both the CNN and EEG-conformer yielded *d*-prime values significantly exceeding chance levels (Fig. 3D; *P* < 0.05, Welch’s *t*-test, compared with randomized datasets). These results demonstrate that ECoG/LFP patterns corresponding to anticipated outcomes associated with emotional valence emerged during CS presentation and could be classified using machine learning. Overall, *d*-prime values did not differ significantly between the CNN and EEG-Conformer models (Fig. 3E; *n* = 5 days; *t*4 = 2.25, *P* = 0.088, Paired *t*-test). To examine the features driving classification, SHAP analysis was applied to the trained EEG-Conformer (Fig. 3F). The absolute SHAP values did not differ substantially across brain regions (*P* = 0.32, Freedman test), suggesting that each region contributed to the classification to a comparable extent. Similarly, we evaluated whether machine learning could classify ECoG/LFP patterns between USr and USp periods, during which the monkey directly received the outcomes (Fig. 3G). In all five recording days, both the CNN and EEG-conformer yielded *d*-prime values significantly exceeding chance levels (Fig. 3H; *P* < 0.05, Welch’s *t*-test, compared with randomized datasets). In addition, *d*-prime values obtained with the EEG-Conformer model were significantly higher than those with the CNN (Fig. 3I; *n* = 5 days; *t*4 = 5.97, *P* = 0.0040, Paired *t*-test). The absolute SHAP values for this classification did not differ substantially across brain regions (Fig. 3J; *P* = 0.55, Freedman test).

**Figure 3.**
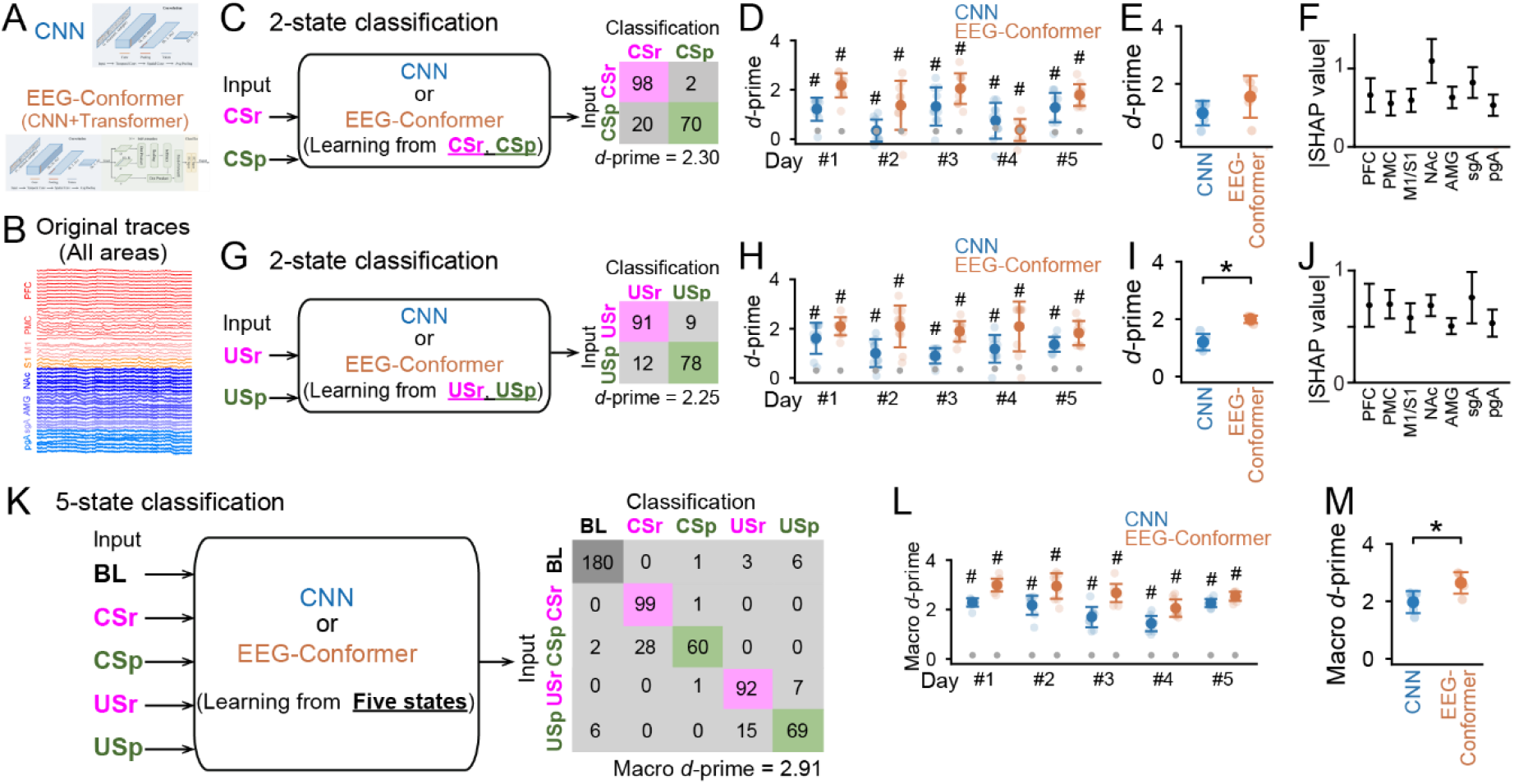
Machine learning-based classification of five states from ECoG/LFP traces. (A, B) Original ECoG/LFP traces from all 50 recording sites were analyzed by the CNN or EEG-Conformer. (C) Classification of CSr and CSp periods. CSr and CSp periods were trained in the CNN or EEG-Conformer with a supervised learning of correct labels, and the other CSr and CSp periods without training were classified as a CSr and CSp period. The right panel shows a matrix representing the number of periods in ten-fold cross-validation from a single recording day, where the values were obtained by summing the test-set predictions across all 10 folds, with the best model in each fold selected based on validation performance and evaluated on the held-out test set. Classification performance (*d*-prime) was defined from each matrix. (D) Comparison of *d*-prime values between CNN and EEG-Conformer in each day. Gray dots represent the upper limit of the 95% critical confidence interval of *d*-prime values computed from randomized datasets. # represents significant differences (*P* < 0.05, Welch’s *t*-test), compared with randomized datasets. (E) Summarized data from the five days shown in D. (F) SHAP values for each brain region derived from EEG-Conformer. *P* = 0.32, Freedman test. (G–J) Same as C–F but for classification of USr and USp periods. \**P* = 0.0039, paired *t*-test (I). *P* = 0.55, Freedman test (J). (K) Classification of five states. Five periods were trained in the CNN or EEG-Conformer, and all remaining periods were classified into one of the five periods. A confusion matrix shows classification of five periods and the macro *d*-prime values averaged from five *d*-prime values from individual periods. (L) Comparison of macro *d*-prime values between CNN and EEG-Conformer in each day. Gray dots represent macro *d*-prime values computed from randomized datasets. # represents significant differences (*P* < 0.05, Welch’s *t*-test), compared with randomized datasets. (M) Summarized data from the five days shown in L. \**P* = 0.0043, paired *t*-test.

Furthermore, we evaluated whether the machine learning models could classify ECoG/LFP patterns across all five states (Baseline, CSr, CSp, USr, and USp), which are considered to differ in both arousal and emotional levels (Fig. 3K). Models were trained simultaneously on all five states and assessed for their ability to correctly identify each state. In this case, classification performance was represented as a 5 × 5 confusion matrix, from which *d*-prime values were derived for each state. Averaged *d*-prime values were computed as a macro *d*-prime value for each dataset. Overall, we confirmed that macro *d*-prime values significantly exceeded chance levels on each day with both models (Fig. 3L; *P* < 0.05, Welch’s *t*-test, compared with randomized datasets), and that those obtained with the EEG-Conformer model were significantly higher than those with the CNN (Fig. 3M; *n* = 5 days; *t*4 = 5.85, *P* = 0.0043, Paired *t*-test). Taken together, both models successfully decoded emotional valence from ECoG/LFP patterns, in the monley with the EEG-Conformer demonstrating consistently superior performance, establishing it as the preferred architecture for valence decoding.

### Classification performance from surface ECoG and intracranial LFP traces

The analyses described so far were performed using ECoG/LFP signals from all recording sites. However, extracellular signals from cortical surface and deep brain regions may carry information of differing quality for estimating emotional valence and arousal level.

We therefore repeated the same analyses separately for each electrode group.

Surface ECoG signals were composed of 25 sites spanning the prefrontal cortex (PFC; 12 sites), premotor cortex (PMC; 6 sites), and primary motor/somatosensory cortex (M1/S1; 7 sites) (Fig. 4A). Even when using the surface ECoG signals alone, *d*-prime values for CSr and CSp classification and USr and USp classification by both the CNN and EEG-Conformer significantly exceeded chance levels in the majority of days (Fig. 4B, top and middle; *P* < 0.05, Welch’s *t*-test, compared with randomized datasets), with no significant difference between the two models (CS: *n* = 5 days, *t*4 = 1.85, *P* = 0.14; US: *t*4 = 2.17, *P* = 0.096, Paired *t*-test). For the 5-state classification, macro *d*-prime values with both models significantly exceeded chance levels in each day (Fig. 4B, bottom; *P* < 0.05, Welch’s *t*-test, compared with randomized datasets), and the EEG-Conformer significantly outperformed the CNN (*t*4 = 7.11, *P* = 0.0021, Paired *t*-test). These results demonstrate that the EEG-Conformer model can discriminate surface ECoG patterns associated with the five states related to different levels of arousal and emotional valence.

**Figure 4.**
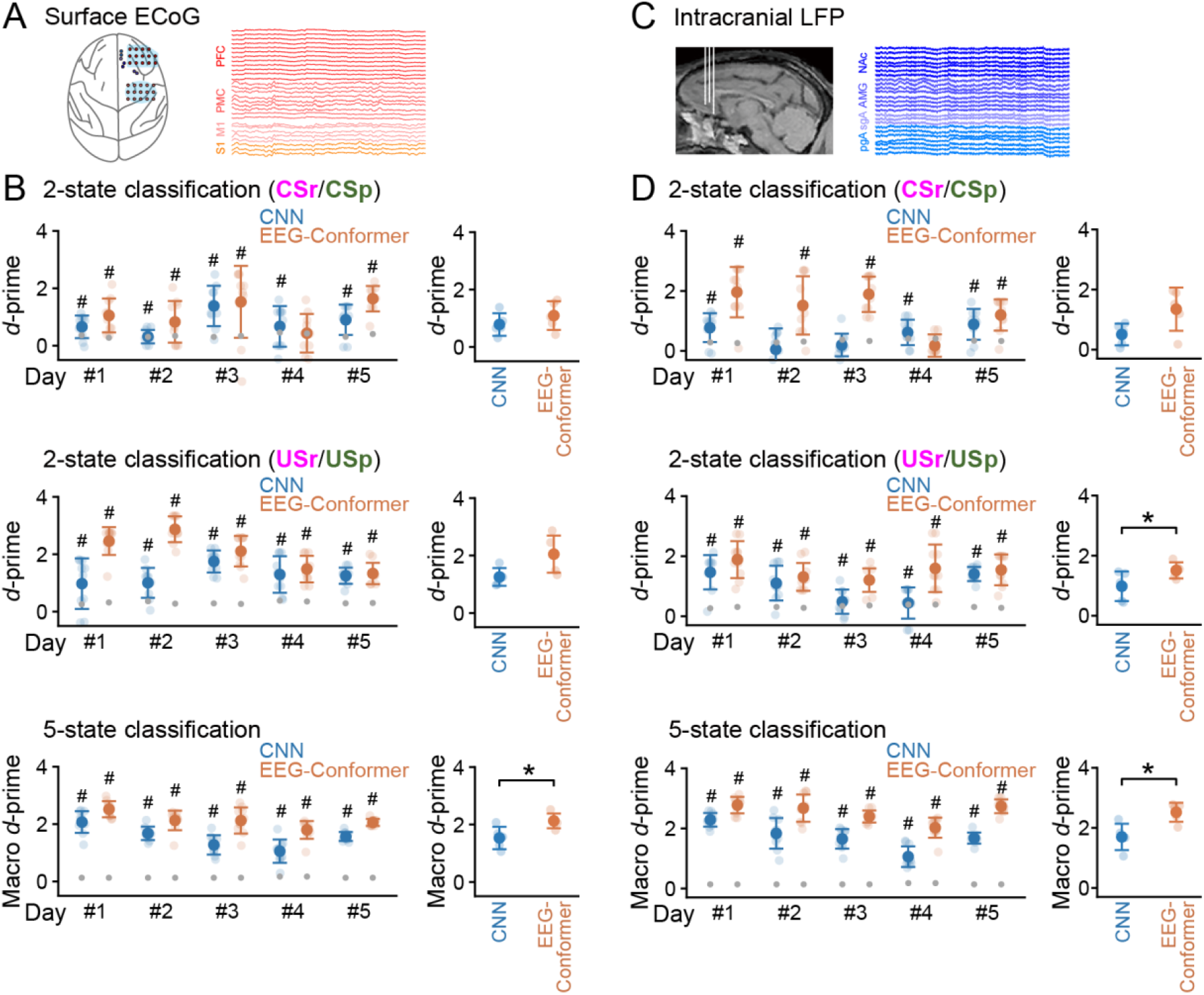
Classification performance when surface ECoG and intracranial LFP traces were analyzed separately. (A) Only surface ECoG traces were analyzed. (B) Comparison of (macro) *d*-prime values between CNN and EEG-Conformer to classify CSr and CSp (top), USr and USp (middle), and five states (bottom). The left panels show *d*-prime values from each day. Gray dots represent the upper limit of the 95% critical confidence interval of *d*-prime values computed from randomized datasets. # represents significant differences (*P* < 0.05, Welch’s *t*-test), compared with randomized datasets. The right panels show summarized data from the five days. \**P* = 0.0021 (bottom), paired *t*-test. (C, D) Same as A and B but when only intracranial LFP traces were analyzed. \**P* = 0.046 (middle) and 0.0011 (bottom), paired *t*-test.

Intracranial LFP signals were composed of 30 sites including the pregenual anterior cingulate cortex (pgACC; 8 sites), subgenual anterior cingulate cortex (sgACC; 7 sites), nucleus accumbens (NAc; 7 sites), and amygdala (AMG; 8 sites) (Fig. 4C). When using the intracranial LFP signals alone, *d*-prime values for CSr and CSp classification by both the CNN and EEG-Conformer significantly exceeded chance levels in the majority of days (Fig. 4D, top; *P* < 0.05, Welch’s *t*-test, compared with randomized datasets), with no significant difference between the two models (*n* = 5 days, *t*4 = 2.13, *P* = 0.10, Paired *t*-test). These *d*-prime values were not significantly different between surface ECoG and intracranial LFP signals (CNN: *n* = 5 days, *t*4 = 1.19, *P* = 0.30; EEG-Conformer: *n* = 5 days, *t*4 = 0.97, *P* = 0.39, Paired *t*-test). For USr and USp classification, *d*-prime values with both models significantly exceeded chance levels in each day (Fig. 4D, middle; *P* < 0.05, Welch’s *t*-test, compared with randomized datasets), and the EEG-Conformer significantly outperformed the CNN (*t*4 = 2.85, *P* = 0.046, Paired *t*-test). These *d*-prime values were not significantly different between surface ECoG and intracranial LFP signals (CNN: *n* = 5 days, *t*4 = 0.83, *P* = 0.45; EEG-Conformer: *n* = 5 days, *t*4 = 1.65, *P* = 0.18, Paired *t*-test). For the 5-state classification, macro *d*-prime values with both models significantly exceeded chance levels in each day (Fig. 4D, bottom; *P* < 0.05, Welch’s *t*-test, compared with randomized datasets), and the EEG-Conformer significantly outperformed the CNN (*t*4 = 8.35, *P* = 0.0011, Paired *t*-test). When using the CNN model, these *d*-prime values were not significantly different between surface ECoG and intracranial LFP signals (*n* = 5 days, *t*4 = 2.74, *P* = 0.052, Paired *t*-test). When using the EEG-Conformer, *d*-prime values were significantly higher from intracranial LFP signals than surface ECoG signals (*n* = 5 days, *t*4 = 4.30, *P* = 0.013, Paired *t*-test). These results suggest that intracranial LFP signals are also discriminable across these emotion-related states, with the EEG-Conformer tending to outperform the CNN.

### Machine learning-based decoding at successive time points

We next applied the EEG-Conformer-based decoder to assess whether distinct emotional states could be resolved across time, tracking classification performance at each successive time point from continuously evolving ECoG/LFP patterns. To enable classification across successive time frames spanning both CS and US epochs within a single trial, the EEG-Conformer was trained on a mixed CS/US dataset with varying CS:US ratios (Fig. 5A). After training on datasets with varying CS:US ratios, classification performance during CS periods or US periods was assessed by computing *d*’CS or *d*’US, respectively (Fig. 5B). When the CS:US ratio was 0:10, meaning that only US periods were included in training, *d*’CS was lowest and *d*’US was highest. As the proportion of CS trials incorporated into training gradually increased, *d*’CS increased progressively while *d*’US decreased correspondingly, such that at a ratio of 10:0, *d*’CS was highest and *d*’US was lowest. These results confirm that classification performance for a given period improves as the proportion of that period’s data in the training set increases. To identify the optimal condition for classifying both CS and US periods, we computed a *J* score as follows (Fig. 5C):

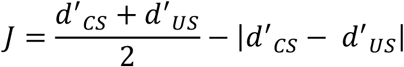

**Figure 5.**
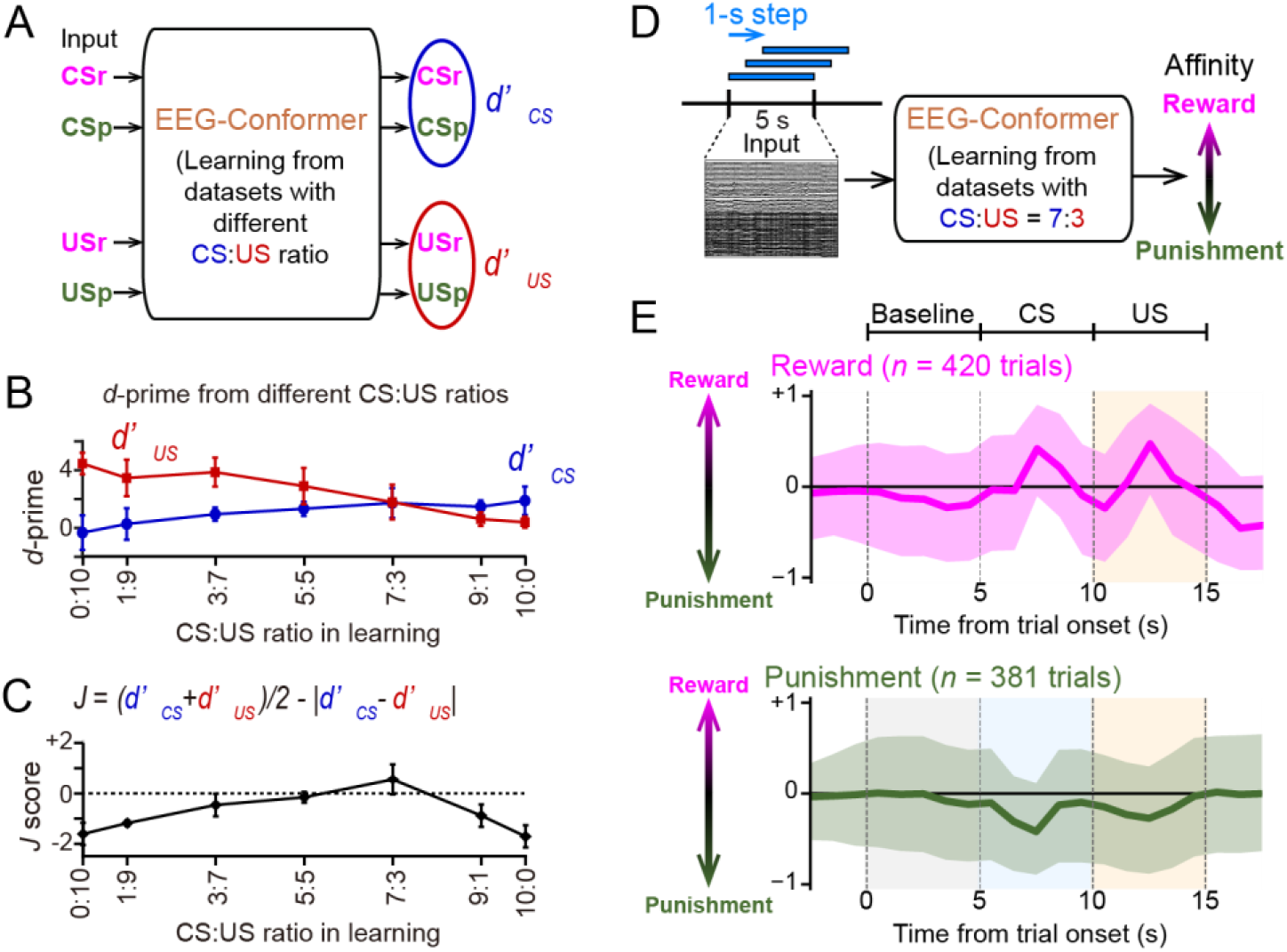
Decoding of emotional valence at continuous time point. (A) The EEG-Conformer was trained on datasets combining CS and US periods at varying CS-to-US ratios, and each CS period and each US period were classified independently, computing *d*’CS and *d*’US, respectively. (B) *d*’CS and *d*’US as a function of CS:US ratios in the training dataset (*n* = 5 days). (C) *J* scores were computed from *d*’CS and *d*’US in B. (D) The EEG-Conformer was trained from datasets with a CS:US ratio of 7:3. ECoG/LFP traces were extracted in 5-s windows advanced in 1-s steps, and all windows not used for training were subjected to classification to estimate affinity to CS/US signals from reward or punishment trials. (E) Averaged temporal changes in the degree of affinity to reward or punishment trials (*n* = 420 and 381 trials, respectively). Lines indicate the median and shaded areas indicate the interquartile range.

We found that the *J* score was maximized at a CS:US ratio of 7:3, indicating that this ratio yielded the most balanced and accurate classification of both periods. For subsequent analyses, we thus used the EEG-Conformer trained on a dataset with the optimal CS:US ratio of 7:3.

In each trial, ECoG/LFP patterns were extracted within a 5-s sliding time window across the 15-s trial duration, and signals from all recording sites were input to the EEG-Conformer to compute affinity scores for reward and punishment trials, output as positive and negative values, respectively (Fig. 5D). The time course of affinity scores for each trial type, averaged across all trials, is summarized in Fig. 5E. In reward trials, affinity scores remained near zero or were weakly negative during the baseline period. Upon CSr presentation, affinity scores gradually shifted in the positive direction, reaching a positive peak approximately 2.5 s after CSr onset. Subsequently, upon USr delivery, affinity scores reached a second positive peak during the 2.5-s reward consumption period. In punishment trials, affinity scores remained near zero or were weakly negative during the baseline period. Upon CSp and USp presentation, responses were diametrically opposite to those in reward trials. Affinity scores gradually shifted in the negative direction, reaching a negative peak approximately 2.5 s after CSp onset. These results demonstrate that the EEG-Conformer model successfully tracked the continuous temporal evolution of emotional states.

## DISCUSSION

In this study, we recorded the ECoG/LFP patterns from superficial and deep brain regions in a monkey that were assumed to be involved in emotion and affect. We first confirmed that overall LFP power in frequency bands below the slow gamma range decreased across most brain regions during USr presentation; however, this change was not sufficiently large to separate USr from USp on a trial-by-trial basis. Both the EEG-Conformer and CNN were applied to raw LFP time-series without conversion to LFP power, both consistently achieving classification performance significantly above chance level, with the EEG-Conformer yielding superior accuracy for both US-type and 5-state classification. This advantage was consistent across both surface ECoG and intracranial LFP signals. Furthermore, by training the EEG-Conformer on a mixed dataset combining CS and US trials (7:3 ratio), we were able to estimate emotion-related changes in brain electrophysiological signals in a temporally continuous manner.

The EEG-Conformer provided a better classification accuracy than CNN, which was consistent with our previous study classifying the neural activity during the abdominal pain or no pain in mice ^8^. The superior performance of the Transformer model is likely attributable to its architectural strength in processing sequential time-series data with complex temporal dependencies, a property that has similarly driven its success in natural language processing. While the attention mechanisms underlying this model do not readily permit identification of the specific features or time points driving classification, our findings demonstrate its robust efficacy in decoding neural signals, including ECoG and LFP recordings, and in resolving intricate ensemble activity patterns associated with emotional processing, stimulus responses, and higher-order cognitive functions. These results suggest that Transformer-based approaches hold considerable promise for real-time neural decoding on a trial-by-trial basis, with broad potential for practical implementation across a variety of brain-machine interface and clinical applications.

It is noteworthy that, by training the EEG-Conformer on a mixed dataset combining CS and US trials, we were able to establish a time-resolved decoder that can detect emotional changes during both the CS and US periods. Because the sensory inputs and motor events that occur during these two periods are quite different, namely, visual stimulation during the CS period, and gustatory stimulation in the mouth accompanied by consummatory movements such as sucking and swallowing the juice, or avoidance movements such as dribbling the saline out of the mouth during the US period, the EEG-Conformer can be considered to have acquired the ability to decode emotion across different sensory and motor contexts. However, to enable the model to acquire this general capability, it was necessary to optimize the composition of the training data by adjusting the ratio of CS to US trials. This finding suggests a practical challenge: to develop a decoder capable of extracting the emotional component across a variety of incidents, the composition of the training dataset must be chosen appropriately. To validate the general utility of this decoder more objectively, it would be useful to examine its behavior in response to novel events that were not included in the training data. This could be achieved by investigating how the decoder behaves when the monkey is presented with novel stimuli that it clearly prefers or finds aversive.

To address the question of which brain regions contained more information related to emotion, we compared the results of spectral analyses across regions, analyzed separately the data from the surface ECoG electrodes and the deep-brain LFP electrodes, and performed SHAP analysis on decoding based on all electrodes. None of these analyses indicated that signals from any specific brain region contained more information than others. This finding is somewhat counterintuitive considering previous reports from single-unit and multi-unit recordings, as well as from functional brain imaging studies using fMRI or PET, which have often identified strong emotion-related signals in regions such as the amygdala and nucleus accumbens. However, in our previous study comparing neural activity between conditions with and without abdominal pain ^8^, we obtained a similar result, namely that signals recorded from electrodes in the amygdala, nucleus accumbens, and other brain regions did not differ greatly in their information content. This may suggest that ECoG and deep LFP signals reflect information distributed over broader regions rather than purely local information at the electrode site. However, other analyses, such as temporal correlation analysis between electrodes or signal source estimation, may help to clarify the localization of the relevant information. Further investigation will therefore be needed. From a practical point of view, the absence of a marked difference in information content between deep LFP signals and surface ECoG signals suggests that the present findings may have meaningful potential for application to scalp EEG recordings in humans.

## Methods

### Subject

One Japanese monkey (Macaca fuscata) (monkey Be: male, 12 years of age, 8.0–9.2 kg, during experiment periods) were used in this study. All procedures were conducted in accordance with the National Institutes of Health (NIH) Guide for the Care and Use of Laboratory Animals and the Tohoku University’s Guidelines for Animal Care and Use. This project was approved by the Center for Laboratory Animal Research of Tohoku University. They were housed individually in cages fulfilling the standards specified in NIH and Tohoku University Guidelines (approximately W750 × D900 × H900 mm) in a room with a 12:12 light-dark cycle (lights on at 08:00). The animal was supplied with water *ad libitum* and fed with monkey chow and fresh vegetables twice daily.

#### Behavioral task

The monkey performed a Pavlovian conditioning task. Conditioned stimuli (CSs) consisted of syringes of two different colors (orange and blue), which were associated with a rewarding and aversive outcomes, respectively. One syringe contained apple juice (unconditioned stimulus reward; USr), whereas the other contained saturated saline (US punishment; USp). In each trial, a single experimenter (S.N.) manually presented one of the colored syringes in front of the monkey for 5 s at a distance of approximately 30 cm from the monkey’s face, serving as the CS (CS reward, CSr, or CS punishment, CSp). The order of CS presentation was pseudo-randomized across trials, such that the number of CSr and CSp trials was balanced within each session and no more than three identical CSs were presented consecutively. Following CS presentation, the experimenter brought the tip of the syringe to the monkey’s mouth, and the liquid was delivered for 5 s as the US. The interval between the end of the US and the onset of the next CS (intertrial interval, ITI) was approximately 15s. The 5-s period preceding CS onset was defined as the baseline (BL). The timing of CS and US presentation was controlled using an LED indicator placed behind the monkey, which served as a timing cue for the experimenter. During the experiment, the monkey was seated in a custom-made primate chair, and its head was noninvasively fixed using a splinting material (Polyflex II, SAKAI Medical Co., Ltd., Japan) customized to conform the lateral aspects of the monkey’s head ^19,20^.

#### Surgery and electrophysiological recording

We used custom-designed electrocorticogram (EcoG) and local field potential (LFP) electrode arrays (Unique Medical Co., Ltd., Japan) for recording neural activity from the brain surface and deep brain structures (Fig 1D). The EcoG electrode array consisted of 14 or 15 platinum plates of 1 mm diameter embedded in a silicon sheet at intervals of 4 mm (Fig. 1D, left). The LFP electrode array consisted of platinum-iridium alloy wires embedded in a Teflon tube (0.3 mm diameter) and have five contacts at intervals of 2 mm (Fig.1D, right).

Preceding the electrode implantation surgery, a structural magnetic resonance and computed tomography images of the monkey’s head was taken under anesthesia induced with a combination of medetomidine (0.04 mg/kg, i.m.), midazolam (0.3 mg/kg, i.m.), and butorphanol (0.4 mg/kg, i.m.). A three-dimensional model of the brain and skull were reconstructed and used for determining the locations of electrode implantation. For the implantation surgery, the animals were anesthetized initially with ketamine (10 mg/kg, i.m.) and xylazine (0.5 mg/kg, i.m), followed by isoflurane (1–2%) inhalation for maintenance. The ECoG electrode arrays were implanted subdurally over the prefrontal (PFC), premotor (PMC), primary motor (MI), and primary somatosensory (SI) cortices in the right hemisphere. The shape of electrode arrays was as shown in Fig. 1D. The LFP electrodes arrays were implanted into the pregenual and subgenual anterior cingulate cortex (pgACC and sgACC), nucleus accumbens (NAc), and amygdala (AMG). Lead lines from the electrodes were connected to connectors (Omnetics Connector Corporation, Minesota, USA). The connectors were set in an acrylic chamber and fixed the posterior part of the skull with dental cement (Super-Bond, Sun Medical, Japan). Electrophysiological signals were amplified and digitalized at 2 kHz using a digital signal processing unit (Cerebus™ system, Blackrock Microsystems, USA) and stored in a computer.

### ECoG/LFP power

To compute the time-frequency representation of the ECoG/LFP power change with time, signals were downsampled to 200 Hz and convolved using a FFT analysis or Morlet’s wavelet family defined by a constant ratio of f0/σf = 5, where f0 represents the frequency of interest and σf represents the bandwidth of the wavelet in the frequency domain chosen to ensure optimal time-frequency localization properties of the wavelet. The frequency bands were defined as follows: delta: 1–4 Hz, theta: 4–8 Hz, alpha: 8–14 Hz, beta: 14–30 Hz, slow gamma: 30–80 Hz, and fast gamma: 80–95 Hz.

For each original trial, baseline, CS, and US epochs were paired, and incomplete triplets or predefined noisy trials were excluded. For each region and frequency band, amplitudes were averaged across electrodes belonging to the same region. To remove trial-wise baseline activity, CS and US amplitudes were corrected by subtracting the amplitude of the corresponding baseline epoch from the same original trial.

Reward–punishment discriminability was evaluated separately for CS and US epochs. For each session, region, and frequency band, signed *d*-prime was calculated as

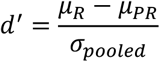

where μR and μP denote the mean baseline-subtracted amplitudes for reward and punishment trials, respectively. The pooled standard deviation was defined as

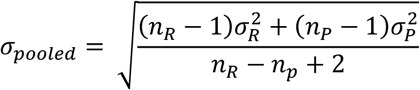

where nR and nP are the numbers of reward and punishment trials, and σR and σP are their standard deviations. Positive *d*-prime values indicate larger reward-related amplitudes than punishment-related amplitudes, whereas negative values indicate larger punishment-related amplitudes.

For group-level visualization and statistics, *d*-prime values were first computed independently for each recording session. The session-wise *d*-prime values were then averaged across sessions for each region, frequency band, and epoch. For color visualization, mean signed *d*-prime values were shown using the same color scale as in the subsequent machine-learning analyses. The white range was defined from a permutation-based null distribution generated by shuffling reward/punishment labels within each session 5,000 times, recomputing session-wise d′ values, and using the across-session mean *d*-prime distribution to set the *p* < 0.001 threshold.

In addition, we defined a session-wise dominant condition based on trial-level comparisons between reward and punishment in Figure 2. For each session, region, frequency band, and epoch, baseline-corrected band-limited power was compared between reward and punishment trials using one-sided Wilcoxon rank-sum tests in both directions. A session was classified as R-dominant or P-dominant only when reward or punishment showed significantly larger power, respectively, with significance defined as *p* < 0.05. The letter displayed in each box indicates R or P only when that condition was dominant in more than half of the recording sessions. If neither condition exceeded this majority criterion, the box was labeled blank. The corresponding Table 1 reports the full counts as R or P (nR:nP), where nR and nP indicate the numbers of R-dominant and P-dominant sessions, respectively.

### Classification based on the CNN

LFP signals from each fixed 5-s trial window were downsampled from 2000 to 200 Hz and used as model input. The resulting channel-by-time matrices were classified using a convolutional neural network (CNN) designed for multichannel LFP decoding. The CNN comprised an initial temporal convolution, a spatial convolution spanning all channels, and additional temporal feature-extraction layers, followed by batch normalization, ELU activation, average pooling, dropout, and a fully connected classifier. Detailed layer specifications are provided in Supplementary Table S1.

For model evaluation, the data were first divided into training and held-out test sets at a ratio of 9:1 using stratified sampling. Stratified 10-fold cross-validation was then performed on the training set, with one fold used for validation in each iteration. In each fold, the best-performing model on the validation set was selected and subsequently evaluated on the common held-out test set. For binary classification tasks, performance was quantified using *d*-prime. For the five-class task, performance was quantified using macro *d*-prime, defined as the mean of one-vs-rest class-wise *d*-prime values.

### Classification based on the EEG-Conformer

The same downsampled 5-s LFP trial windows were also analyzed using an EEG-Conformer implemented in PyTorch. The model consisted of convolution-based patch embedding, a transformer encoder, and a classification head. The embedding dimension was 40, the encoder depth was 1, and the learning rate was set to 2 × 10^-4^. Performance was evaluated using the same held-out test set and stratified 10-fold cross-validation procedure described above. For binary tasks, performance was quantified using *d*-prime, whereas for the five-class task it was quantified using macro *d*-prime. Detailed layer specifications are provided in Supplementary Table S2.

### Evaluation of classification performance based on d-prime values

In the classification analyses, *d*-prime values were employed as a measure to assess how accurately our analysis could discriminate between the two conditions. The numbers of hit bins (*n*hit, where both the prediction and the real data were post-bins), miss bins (*n*miss, where the prediction was a pre-bin, but the real data were post-bin), false alarm bins (*n*FA, where the prediction was a post-bin, but the real data were pre-bin), and correct rejection bins (*n*CR, where both the prediction and the real data were pre-bins) were computed. The hit rate (*dHit* = *n*hit/(*n*hit+*n*miss)) and false alarm rate (*pFA* = *n*FA/(*n*FA+*n*CR)) were further computed, while the *d*-prime value was calculated as *Z*(*dHit*) – *Z*(*pFA*).

### Statistics

All data are presented as the mean ± SEM, and were analyzed using MATLAB. Normally distributed data are displayed as the sample mean and SEM with individual data points. Comparisons of two-sample data were performed using paired or Student’s *t*-tests. Multiple group comparisons were performed using Tukey’s test after ANOVA. The null hypothesis was rejected at *P* < 0.05 level.

## ACNOWLEDGEMENT

This work was supported by a grant (JPMJMS2292; JPMJKP25Y8) from the Japan Science and Technology Agency (JST) to KI. Tsutsui, and T. Sasaki; and a grant (JPMJCR21P1) from the JST and KAKENHI (21H05243; 23K26159, 24K21735) from the Japan Society for the Promotion of Science (JSPS) to T. Sasaki; and a KAKENHI (22K07316) from JSPS to H. Watanabe.

## AUTHOR CONTRIBUTIONS

S.N. and K.T. designed the study. S.N. acquired the electrophysiological data. T.X., W.H., and T.S. performed the analyses and prepared the figures. T.S. and K.T. supervised the project and wrote the main manuscript text, and all the authors reviewed the main manuscript text.

## DATA AVAILABILITY STATEMENT

The data that support the findings of this study are available from the corresponding authors upon reasonable request.

---

**Supplementary Table S1.** The Architecture of the CNN.

| Block | Layer | Parameters | Output |
| --- | --- | --- | --- |
| Input | Input | $1 \times C \times 1000$ | $1 \times C \times 1000$ |
| Temporal feature extraction | Conv2d +<br>BatchNorm + ELU | 16 filters, kernel (1, 25) | $16 \times C \times 1000$ |
| Spatial feature extraction | Conv2d +<br>BatchNorm + ELU | 32 filters, kernel (C, 1) | $32 \times 1 \times 1000$ |
| Temporal downsampling | Average pooling +<br>Dropout | pool (1, 4), stride (1, 4),<br>dropout 0.5 | $32 \times 1 \times 250$ |
| | Average pooling +<br>Dropout | pool (1, 4), stride (1, 4),<br>dropout 0.5 | $64 \times 1 \times 62$ |
| Feature extraction | Conv2d +<br>BatchNorm + ELU | 64 filters, kernel (1, 15) | $64 \times 1 \times 250$ |
| | Conv2d +<br>BatchNorm + ELU | 64 filters, kernel (1, 9) | $64 \times 1 \times 62$ |
| Global pooling | Adaptive average pooling | output size (1, 1) | $64 \times 1 \times 1$ |
| Classifier | Fully connected +<br>ELU + Dropout | $64 \rightarrow 128$, dropout 0.5 | 128 |
| Output | Fully connected | $128 \rightarrow \text{num\_classes}$ | num_classes |

**Supplementary Table S2.**
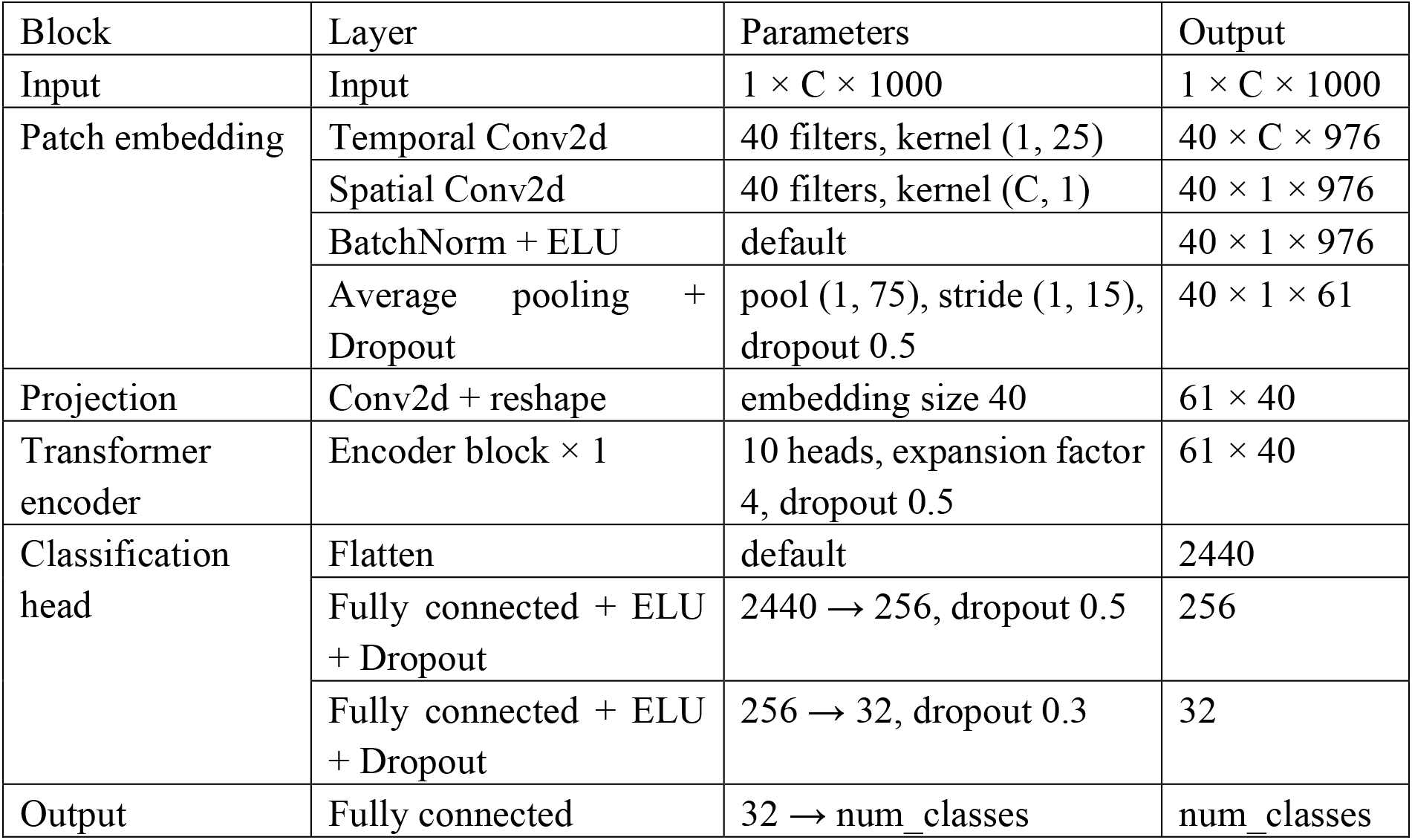
The Architecture of the EEG-Conformer.

